# Stress Tolerance of Multiple *Salmonella enterica* Strains Associated with Foodborne Outbreaks

**DOI:** 10.1101/2025.04.23.649775

**Authors:** A. Martin, C. Patch, V. Vanarsdall, R. Pham, G. Whitney, S. Markus, A. Lunna, A.J. Etter

**Affiliations:** Department of Nutrition and Food Sciences, University of Vermont. Burlington, Vermont

**Author notes:** These authors contributed equally to this work (A. Martin, C. Patch).

**Keywords:** Salmonella, stress tolerance, foodborne outbreak

## Abstract

*Salmonella enterica* is a foodborne pathogen commonly found in food processing environments. While control methods such as heat treatment and sanitizers are often used, *S. enterica* has evolved strategies for survival and persistence to overcome pathogen control. This study assessed 43 outbreak-associated (OA) and non-outbreak associated (NOA) *S. enterica* isolates from serovars Enteritidis, Heidelberg, Newport, Typhimurium, and monophasic Typhimurium (I 4,[5],12:i:-) for enhanced stress tolerance. Heat shock at 56° C, minimum inhibitory concentrations (MICs) for sanitizers sodium hypochlorite (NaOCl) and peracetic acid (PAA), and crystal violet microtiter assays were used to evaluate heat tolerance, sanitizer tolerance, and attachment capacity, respectively. Most isolates (n =34/43) carried at least one antimicrobial resistance gene, and nearly half (n =21/43) displayed genotypic and/or phenotypic resistance to ampicillin, ciprofloxacin, or ceftriaxone. Most isolates carried genes conferring resistance to gold (n =43/43) and arsenic (n= 41/43), and tolerance to mercury, copper, and silver was common among monophasic Typhimurium and Heidelberg isolates. Efflux pump *qacEdelta1* was detected among eight Heidelberg isolates. We found enhanced stress tolerance (i.e. an unusually high ability to survive and adapt to various environmental stresses) to sanitizers and enhanced attachment capacity, indicating biofilm formation. Isolates evaluated for heat tolerance survived at least 15 min at 56° C and three survived >60 min. Overall, we found evidence of enhanced tolerance to individual stresses across both OA and NOA *S. enterica.* There were no strong patterns based upon serovar or OA/NOA status; however, we did find that specific enhanced stress tolerance profiles may have contributed to outbreak characteristics.

## Introduction

Non-typhoidal *S. enterica* (NTS) is a Gram-negative foodborne pathogen commonly found in poultry, eggs, pork, beef, nuts, and produce [1]. NTS is responsible for numerous foodborne outbreaks annually in the United States and is a common contaminant in processing facilities [2]. Salmonellosis also carries a high economic burden, with the loss of over $3.3 billion annually in healthcare costs, lost productivity, and mortality in the U.S. [3]. NTS can exhibit high rates of antimicrobial resistance (AMR), which is cited by the Centers for Disease Control and Prevention (CDC) as a serious public health concern [4]. More specifically, fluoroquinolone resistant NTS is considered a “high concern” pathogen by the World Health Organization [5].

To limit bacterial growth and prevent foodborne outbreaks, various pathogen control methods may be used in the food processing industry, including temperature control (e.g., heat shock) and sanitizers such as sodium hypochlorite (NaOCl) or peracetic acid (PAA) [6–9]. Often, multiple techniques are used in combination to prevent the proliferation of pathogenic microorganisms [10]. Heat treatment is commonly used and highly effective [11], as *S. enterica* can grow on foods held between 4° C and 60° C (40-140° F; i.e. the “temperature danger zone”) [12]. In poultry processing, scalding is commonly used, either by steam-spraying or immersion scalds, with temperature and duration varying [13]. Current FSIS guidelines recommend 30-75 seconds of exposure at 59-64° C for hard scalds and 90-120 seconds at 51-54° C for soft scalds for chickens, while turkey is typically scalded for 50-125 seconds at 59-63° C [14]. Hard scalds are more common in poultry processing than soft scalds [14]. In pork processing, scalding for 8 minutes at 60-62° C, hot water decontamination for 12-15 seconds at 80° C, and steam pasteurization at 70° C are commonly used [15]. Temperature controls are also employed in peanut and tree nut processing to reduce *S. enterica* [16–18]. Almonds should be dry roasted at 120-148° C for 9 min to 3 h, oil roasted at 126° C for 2 minutes or blanched at 80-90° C [16–18]. Dry roasting peanuts at 154° C for 15 min and oil roasting at 150° C for 1.5 min have been sufficient in reducing *Salmonella* by over 5.4 log CFU/g and 6.0 log CFU/g, respectively [19].

*S. enterica* has developed various mechanisms for survival and persistence in the food processing industry [11, 20], including sanitizer, heat tolerance, and biofilm formation [21]. Exposure to one stressor can lead to cross-tolerance to other stressors [21], further enhancing *S. enterica*’s ability to survive and persist. For instance, heat tolerance in *S. enterica* is influenced by factors such as pre-exposure to stress and starvation, growth phase during heat shock, and expression of heat shock proteins [22–25]. Additionally, thermal resistance mechanisms can play a role in modulating virulence [22]. Repeated exposure to antimicrobials, such as sanitizers, can lead to elevated minimum inhibitory concentration (MIC), or the level of sanitizer required to inhibit the growth of or kill the bacteria; however, development of true resistance is rare [26, 27]. *S. enterica* may also form biofilms, which are difficult to eliminate and provide protection against sanitizers, desiccation, antibiotics, and some host defenses [28–31]. These biofilms often form on untreated or mechanically sanded steel surfaces, even when dry and lacking a steady nutrient source [32]. Biofilms are an especially critical target, as an estimated 80% of all U.S. bacterial infections are linked to foodborne pathogens residing in biofilms [31].

Previous research by Etter et al., 2019 highlighted that six *Salmonella* Heidelberg isolates from a 2013-2014 chicken poultry outbreak displayed enhanced heat tolerance and attachment capacity under stressful conditions [23]. This research laid the groundwork for this study, which aims to further investigate enhanced stress tolerances, variation by serovar, and overall mechanisms that may contribute to outbreak characteristics across multiple *S. enterica* serovars and outbreaks. Further, understanding strain variation can improve processing techniques and consumer risk modeling to reduce foodborne illness [11]. The objectives of this research were to characterize (i) AMR and stress associated genes, (ii) sanitizer tolerance, (iii) heat tolerance, and (iv) attachment capacity of *S. enterica* isolates of various serovars to understand intrinsic characteristics contributing to differences between serovars and isolates associated with historically relevant outbreaks, as well as non-outbreak associated isolates.

## Methods

### Acquisition of Culture Collection

48 strains were included in this study (**Table 1**); 25 were obtained from the USDA-ARS Culture Collection (NRRL), six from the Food and Drug Administration (FDA), three from the New York State Food Laboratory, eight from the Cornell Food Safety Lab, and one strain was ordered from the American Type Culture Collection (ATCC).

**Table 1:**
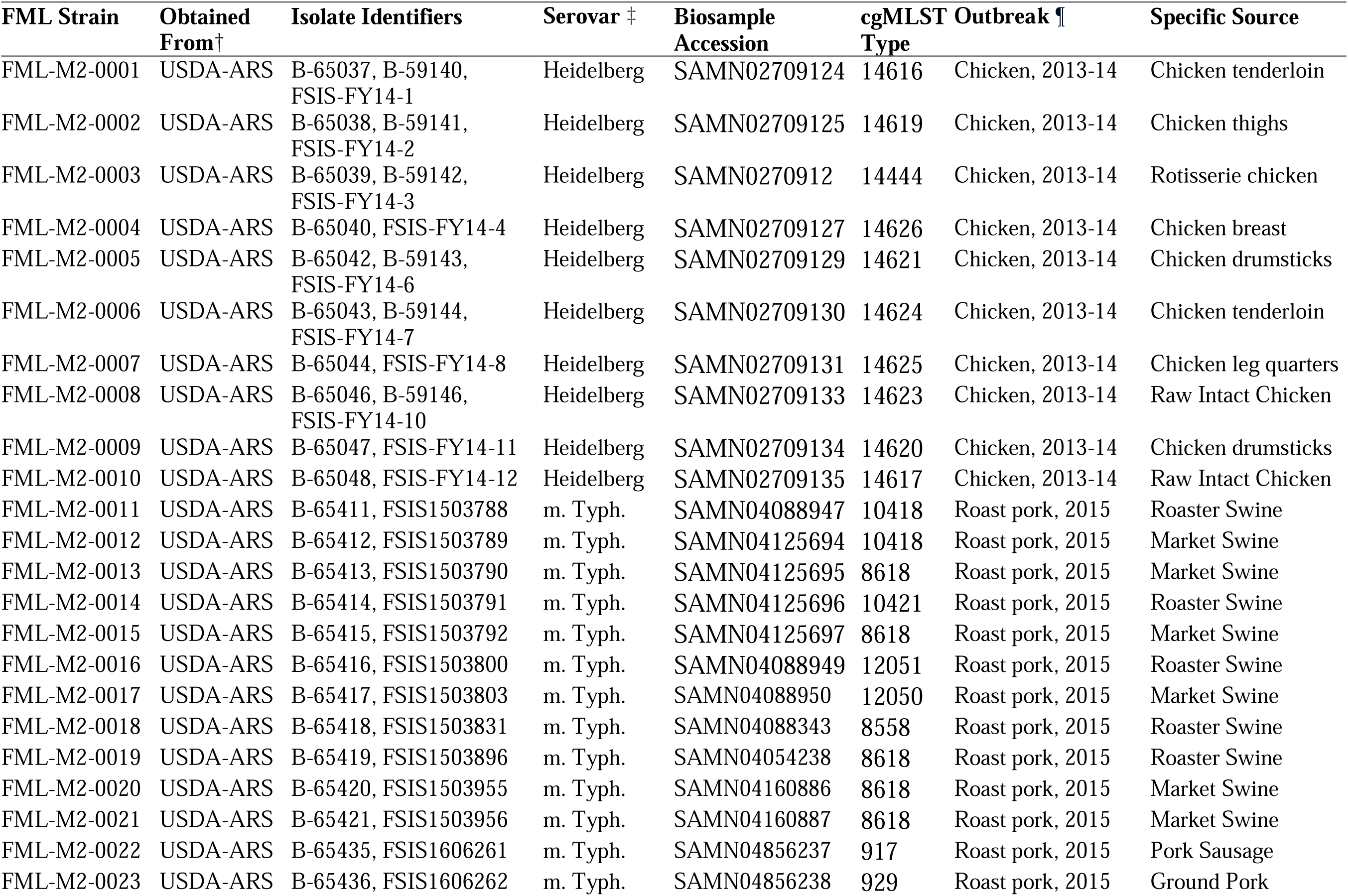
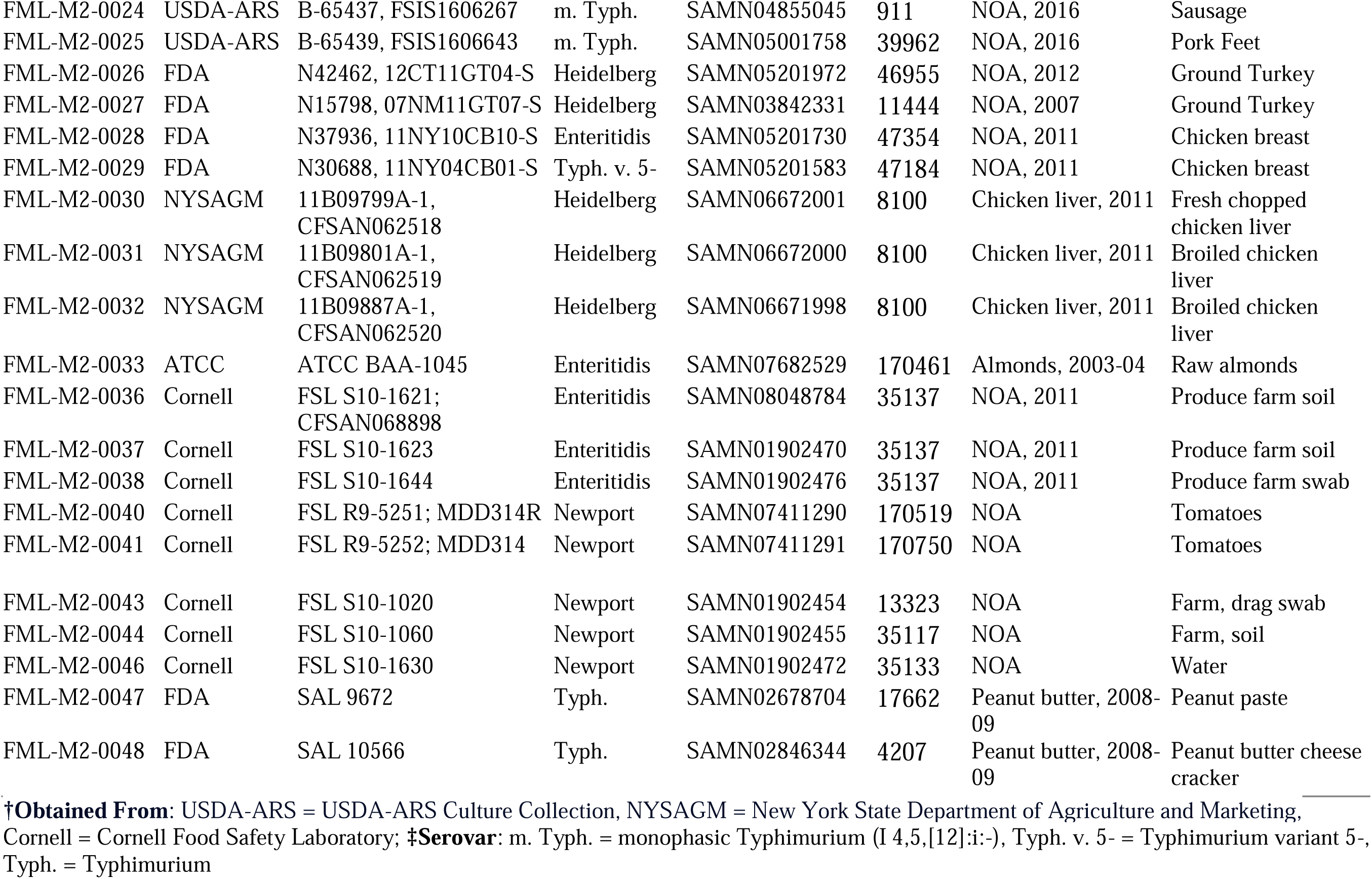
Characteristics of strains used in this study.

| FML Strain | Obtained From† | Isolate Identifiers | Serovar ‡ | Biosample Accession | cgMLST Type | Outbreak ¶ | Specific Source |
| --- | --- | --- | --- | --- | --- | --- | --- |
| FML-M2-0001 | USDA-ARS | B-65037, B-59140, FSIS-FY14-1 | Heidelberg | SAMN02709124 | 14616 | Chicken, 2013-14 | Chicken tenderloin |
| FML-M2-0002 | USDA-ARS | B-65038, B-59141, FSIS-FY14-2 | Heidelberg | SAMN02709125 | 14619 | Chicken, 2013-14 | Chicken thighs |
| FML-M2-0003 | USDA-ARS | B-65039, B-59142, FSIS-FY14-3 | Heidelberg | SAMN0270912 | 14444 | Chicken, 2013-14 | Rotisserie chicken |
| FML-M2-0004 | USDA-ARS | B-65040, FSIS-FY14-4 | Heidelberg | SAMN02709127 | 14626 | Chicken, 2013-14 | Chicken breast |
| FML-M2-0005 | USDA-ARS | B-65042, B-59143, FSIS-FY14-6 | Heidelberg | SAMN02709129 | 14621 | Chicken, 2013-14 | Chicken drumsticks |
| FML-M2-0006 | USDA-ARS | B-65043, B-59144, FSIS-FY14-7 | Heidelberg | SAMN02709130 | 14624 | Chicken, 2013-14 | Chicken tenderloin |
| FML-M2-0007 | USDA-ARS | B-65044, FSIS-FY14-8 | Heidelberg | SAMN02709131 | 14625 | Chicken, 2013-14 | Chicken leg quarters |
| FML-M2-0008 | USDA-ARS | B-65046, B-59146, FSIS-FY14-10 | Heidelberg | SAMN02709133 | 14623 | Chicken, 2013-14 | Raw Intact Chicken |
| FML-M2-0009 | USDA-ARS | B-65047, FSIS-FY14-11 | Heidelberg | SAMN02709134 | 14620 | Chicken, 2013-14 | Chicken drumsticks |
| FML-M2-0010 | USDA-ARS | B-65048, FSIS-FY14-12 | Heidelberg | SAMN02709135 | 14617 | Chicken, 2013-14 | Raw Intact Chicken |
| FML-M2-0011 | USDA-ARS | B-65411, FSIS1503788 | m. Typh. | SAMN04088947 | 10418 | Roast pork, 2015 | Roaster Swine |
| FML-M2-0012 | USDA-ARS | B-65412, FSIS1503789 | m. Typh. | SAMN04125694 | 10418 | Roast pork, 2015 | Market Swine |
| FML-M2-0013 | USDA-ARS | B-65413, FSIS1503790 | m. Typh. | SAMN04125695 | 8618 | Roast pork, 2015 | Market Swine |
| FML-M2-0014 | USDA-ARS | B-65414, FSIS1503791 | m. Typh. | SAMN04125696 | 10421 | Roast pork, 2015 | Roaster Swine |
| FML-M2-0015 | USDA-ARS | B-65415, FSIS1503792 | m. Typh. | SAMN04125697 | 8618 | Roast pork, 2015 | Market Swine |
| FML-M2-0016 | USDA-ARS | B-65416, FSIS1503800 | m. Typh. | SAMN04088949 | 12051 | Roast pork, 2015 | Roaster Swine |
| FML-M2-0017 | USDA-ARS | B-65417, FSIS1503803 | m. Typh. | SAMN04088950 | 12050 | Roast pork, 2015 | Market Swine |
| FML-M2-0018 | USDA-ARS | B-65418, FSIS1503831 | m. Typh. | SAMN04088343 | 8558 | Roast pork, 2015 | Roaster Swine |
| FML-M2-0019 | USDA-ARS | B-65419, FSIS1503896 | m. Typh. | SAMN04054238 | 8618 | Roast pork, 2015 | Roaster Swine |
| FML-M2-0020 | USDA-ARS | B-65420, FSIS1503955 | m. Typh. | SAMN04160886 | 8618 | Roast pork, 2015 | Market Swine |
| FML-M2-0021 | USDA-ARS | B-65421, FSIS1503956 | m. Typh. | SAMN04160887 | 8618 | Roast pork, 2015 | Market Swine |
| FML-M2-0022 | USDA-ARS | B-65435, FSIS1606261 | m. Typh. | SAMN04856237 | 917 | Roast pork, 2015 | Pork Sausage |
| FML-M2-0023 | USDA-ARS | B-65436, FSIS1606262 | m. Typh. | SAMN04856238 | 929 | Roast pork, 2015 | Ground Pork |
| FML-M2-0024 | USDA-ARS | B-65437, FSIS1606267 | m. Typh. | SAMN04855045 | 911 | NOA, 2016 | Sausage |
| FML-M2-0025 | USDA-ARS | B-65439, FSIS1606643 | m. Typh. | SAMN05001758 | 39962 | NOA, 2016 | Pork Feet |
| FML-M2-0026 | FDA | N42462, 12CT11GT04-S | Heidelberg | SAMN05201972 | 46955 | NOA, 2012 | Ground Turkey |
| FML-M2-0027 | FDA | N15798, 07NM11GT07-S | Heidelberg | SAMN03842331 | 11444 | NOA, 2007 | Ground Turkey |
| FML-M2-0028 | FDA | N37936, 11NY10CB10-S | Enteritidis | SAMN05201730 | 47354 | NOA, 2011 | Chicken breast |
| FML-M2-0029 | FDA | N30688, 11NY04CB01-S | Typh. v. 5- | SAMN05201583 | 47184 | NOA, 2011 | Chicken breast |
| FML-M2-0030 | NYSAGM | 11B09799A-1,<br>CFSAN062518 | Heidelberg | SAMN06672001 | 8100 | Chicken liver, 2011 | Fresh chopped<br>chicken liver |
| FML-M2-0031 | NYSAGM | 11B09801A-1,<br>CFSAN062519 | Heidelberg | SAMN06672000 | 8100 | Chicken liver, 2011 | Broiled chicken<br>liver |
| FML-M2-0032 | NYSAGM | 11B09887A-1,<br>CFSAN062520 | Heidelberg | SAMN06671998 | 8100 | Chicken liver, 2011 | Broiled chicken<br>liver |
| FML-M2-0033 | ATCC | ATCC BAA-1045 | Enteritidis | SAMN07682529 | 170461 | Almonds, 2003-04 | Raw almonds |
| FML-M2-0036 | Cornell | FSL S10-1621;<br>CFSAN068898 | Enteritidis | SAMN08048784 | 35137 | NOA, 2011 | Produce farm soil |
| FML-M2-0037 | Cornell | FSL S10-1623 | Enteritidis | SAMN01902470 | 35137 | NOA, 2011 | Produce farm soil |
| FML-M2-0038 | Cornell | FSL S10-1644 | Enteritidis | SAMN01902476 | 35137 | NOA, 2011 | Produce farm swab |
| FML-M2-0040 | Cornell | FSL R9-5251; MDD314R | Newport | SAMN07411290 | 170519 | NOA | Tomatoes |
| FML-M2-0041 | Cornell | FSL R9-5252; MDD314 | Newport | SAMN07411291 | 170750 | NOA | Tomatoes |
| FML-M2-0043 | Cornell | FSL S10-1020 | Newport | SAMN01902454 | 13323 | NOA | Farm, drag swab |
| FML-M2-0044 | Cornell | FSL S10-1060 | Newport | SAMN01902455 | 35117 | NOA | Farm, soil |
| FML-M2-0046 | Cornell | FSL S10-1630 | Newport | SAMN01902472 | 35133 | NOA | Water |
| FML-M2-0047 | FDA | SAL 9672 | Typh. | SAMN02678704 | 17662 | Peanut butter, 2008-09 | Peanut paste |
| FML-M2-0048 | FDA | SAL 10566 | Typh. | SAMN02846344 | 4207 | Peanut butter, 2008-09 | Peanut butter cheese<br>cracker |

†Obtained From: USDA-ARS = USDA-ARS Culture Collection, NYSAGM = New York State Department of Agriculture and Marketing, Cornell = Cornell Food Safety Laboratory; ‡Serovar: m. Typh. = monophasic Typhimurium (I 4,5,[12]:i:-), Typh. v. 5- = Typhimurium variant 5-, Typh. = Typhimurium

Upon acquisition, cultures were transferred into sterile trypticase soy broth (TSB; BD, Franklin Lakes, NJ) and incubated (37°C, 200 rotations per minute (rpm), 24 hours), then streaked out for isolation onto trypticase soy agar (TSA; BD, Franklin Lakes, NJ) and incubated (37°C, 24 hours). Isolates were stored in sterile cryovials in 25% glycerol at -80°C as a working stock for future use.

### Genomic Characteristics

Antimicrobial resistance genes, stress tolerance genes, and heavy metal resistance genes for each isolate were extracted from the NCBI Isolates Browser platform, which detected these genes using AMRFinderPlus (v3.8.4) [33]. A single nucleotide polymorphism (SNP)-based phylogenetic tree was created to assess isolate relationships using CSIphylogeny (v1.4) [34].

### Sanitizer Tolerance

Minimum Inhibitory Concentrations (MICs) for sodium hypochlorite (NaOCl) and peroxyacetic acid (PAA) sanitizer tolerances were determined as follows, using a procedure adapted from methods previously described [23, 35].

#### Preparation of Cultures

Isolates were recovered from working stock solutions; approximately 15-25 isolated colonies per strain were selected via sterile swab and suspended in 3 mL of TSB. Optical density at 600nm (OD_600_) was read and adjusted to 0.600 – 0.800. Adjusted culture was diluted 1:100 into 1 mL of either 2X TSB or 1/10X TSB to achieve a final concentration of 1/20X (nutrient depletion) or 1X (nutrient abundance) upon sanitizer addition. An NaOCl solution (4.275% NaOCl; Clorox, Oakland, CA) was prepared to achieve a concentration of 400 ppm, then serially diluted into phosphate buffered saline (PBS) to 6.25-200 ppm (0.000625-0.02%) upon addition of bacterial culture. A PAA solution (Inspexx^™^ 250, Saint Paul, MN) was prepared to a concentration of 400 ppm and serially diluted 1:1 into PBS to 25-200 ppm (0.0025-0.02%) upon addition of bacterial cultures.

#### Plate Inoculation and Incubation

Polystyrene microtiter plates were prepared at 1/20X and 1X conditions per strain in triplicate. Plates were read immediately following inoculations to determine a baseline OD_600_ and read again at 24 hours to measure growth at room temperature (22°C), where the MIC was the concentration of sanitizer at which no growth occurred in the wells.

### Heat Shock

Growth curves and heat shock assays were performed to determine heat tolerance using procedures adapted from methods previously described by Etter et al. [23]. Isolates were recovered from the working stock solutions, inoculated on TSA, and incubated (37° C, 24 hours). A single colony was inoculated into 10 mL TSB and incubated (37° C, 200 rpm, 16 hours) before serial dilution in duplicate by a factor of 10^-5^ and incubation (37° C, 200 rpm, 8 hours).

For growth curves, 1 mL aliquots were collected each hour for eight hours and serially diluted into PBS. Dilutions were spread-plated onto TSA in duplicate and incubated (37° C, 36 hours). For heat shock, 1 mL aliquots were transferred to preheated 56° C water baths (hard scald temperature [23, 36]) for heat shock, staggering incubation by 3 minutes per strain. At 0, 3, 6, 9, 15, 30, 45, and 60 minutes, 1 mL aliquots were removed and serially diluted into PBS. Dilutions were immediately pour-plated with 10 mL liquid TSA in duplicate. At 15, 30, 45, and 60 minutes, non-diluted cultures were pour-plated in 100 µL aliquots in duplicate and 250 µL aliquots in quadruplicate to capture low titer cultures, and plates were incubated (37° C, 36 hours). Colonies were counted using a countable range of 30-300 colonies/plate, all counts from 250 µL plates were used, and the limit of detection was 1 CFU/mL.

### Biofilm Attachment Assays

Attachment to polystyrene plates was evaluated using a crystal violet staining procedure adapted from methods previously described [23, 35]. Polystyrene 96-well plates were prepared under four conditions: 1/20X TSB, 4°C (nutrient depletion, refrigeration); 1/20X TSB, RT (22°C) (nutrient depletion, room temperature); 1X TSB, 4°C (nutrient abundance, refrigeration); and 1X TSB, RT (nutrient abundance, room temperature) in triplicate per strain. Absorbance at 600nm was recorded to determine crystal violet (CV) retention as an approximation of cell density (biomass) with an Epoch microplate spectrophotometer at 24, 72, and 120 hours (Agilent Technologies, Santa Clara, CA). Growth was not confirmed via enumeration. All CV attachment assays were repeated in their entirety in triplicate.

### Statistical Methods

All significant differences for the crystal violet attachment assays, the minimum inhibitory concentration assays, and the heat shock assays were assessed using Analysis of Variance (ANOVA), followed by Tukey’s honest significance test in R v4.2.1 [37].

## Results and Discussion

### Genotypic Profiles

A SNP-based phylogenetic tree of all isolates, aligned against reference *Salmonella* Heidelberg SL476 is available in **figure S1**. All *Salmonella* Heidelberg genomes, both outbreak and non-outbreak associated isolates, were within 130 SNPs of each other. Isolates from the Kosher Broiled chicken liver [38, 39] outbreak were zero SNPs different from each other, while isolates from the 2013-2014 poultry outbreak, which involved six strains [40], were 1-122 SNPs different. Monophasic *Salmonella* isolates were 1-102 SNPs different, including the NOA isolate. The two *Salmonella* Typhimurium isolates from the 2008-2009 Peanut Butter outbreak [41] were 73 SNPs different, despite sharing the same two-enzyme PFGE patterns (JPXX01.0459/JPXA26.0462). The *Salmonella* Enteritidis almond outbreak isolate [42] was the most different from its paired NOA isolates; while the NOA isolates were 44-45 SNPs different from each other, they were 2440-2449 SNPs different from the OA isolate. This may have been due to source; the NOA isolates were from produce and soil.

#### Antimicrobial Resistance Genes

Antimicrobial resistance is a concern for *S. enterica* outbreaks as resistance to medically relevant antibiotics (defined as ampicillin, ciprofloxacin, azithromycin, ceftriaxone, and amoxicillin can complicate the treatment of severe clinical cases) and AMR genes often travels on plasmids which can carry virulence, AMR, and sanitizer efflux genes [4, 43–46]. We were interested in whether AMR might be connected to stress tolerance in outbreak-associated *S. enterica*

Most isolates (n = 34/43) carried at least one gene conferring AMR (**Figure 1**). Nearly half (n =21/43) harbored genes associated with resistance to at least one medically important antibiotic. Three Heidelberg isolates from the 2013-2014 chicken outbreak and all monophasic Typhimurium isolates from the 2015 roast pork outbreak carried *blaTEM-1,* conferring ampicillin resistance. Additionally, *blaTEM-1* was detected in one NOA isolate each from serovars Heidelberg, monophasic Typhimurium, and Enteritidis. Newport isolate FML-M2-0046 (NOA, water), contained *blaCMY-2*, linked to third-generation cephalosporin resistance [47, 48]. NOA monophasic Typhimurium isolate FML-M2-0025 (NOA, pork feet) contained *blaSHV-12* (linked to ampicillin and cephalosporin resistance) [49] and *qnrB19* (intermediate quinolone resistance, defined as 0.12–0.5 mg/L [50]); it was the only isolate to carry either gene. Phenotypic resistance data (**Table S1**, available for a limited number of isolates) showed that monophasic Typhimurium isolate FML-M2-0025 (NOA, pork feet) also exhibited intermediate ciprofloxacin resistance and Enteritidis isolate FML-M2-0028 (NOA, chicken breast) exhibited intermediate resistance to amoxicillin/clavulanic acid. As for genotypic resistance among NOA isolates, half (n =7/14) contained no AMR genes, while AMR genes were particularly prevalent among Heidelberg isolate FML-M2-0026 (NOA, ground turkey), Enteritidis isolate FML-M2-0028 (NOA, chicken breast), and Newport isolate FML-M2-0046 (NOA, water).

**Figure 1:**
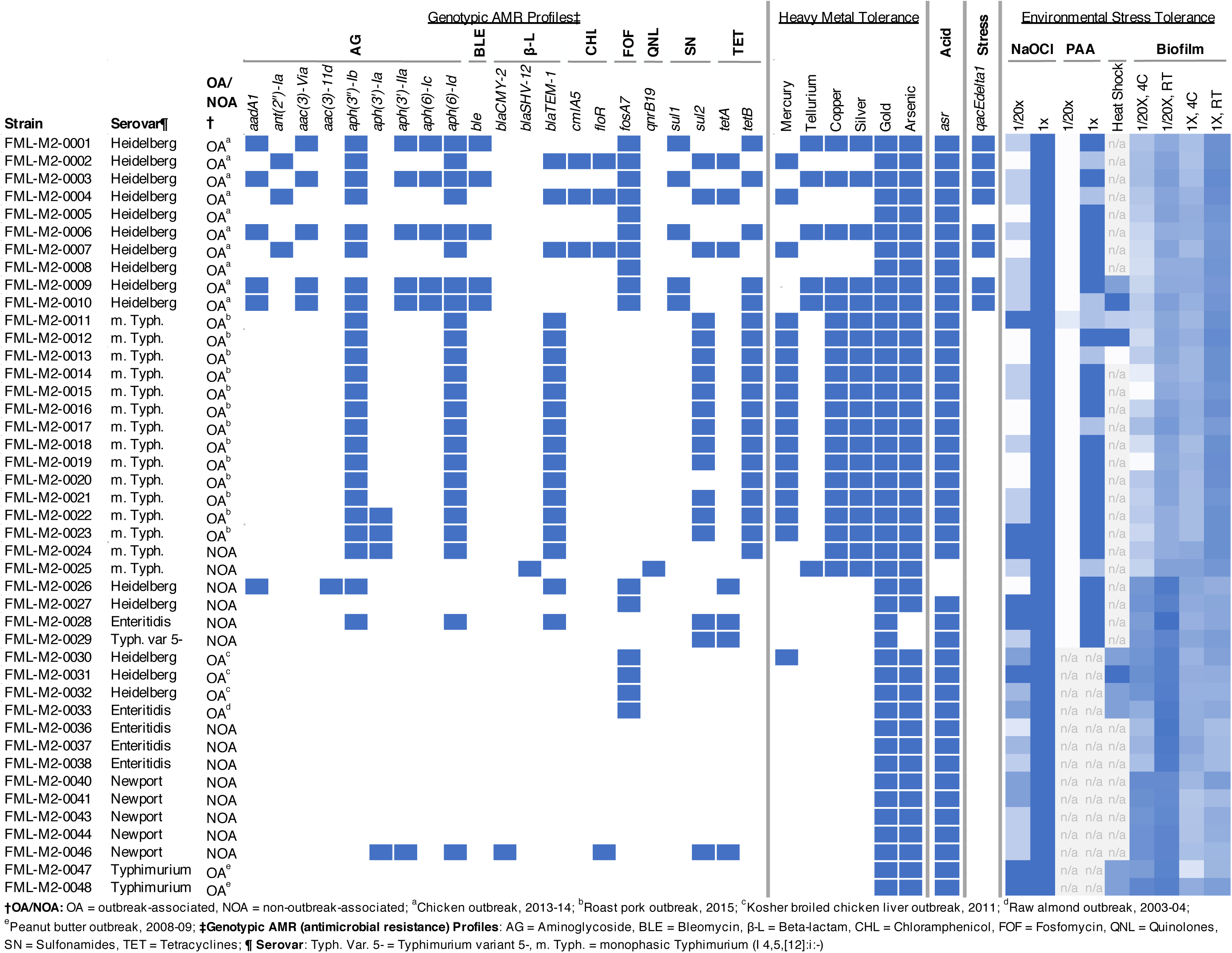
Heatmap of isolate antimicrobial resistance profiles, relevant genotypic characteristics, and results of environmental stress tolerance experiments. Heatmap values for environmental stress tolerance are based upon MIC for sanitizers, absorption values for biofilm formation (OD_600_), and the timepoint to which isolates survived 56°C heat shock (30, 45, 60 or >60 minutes).

Efflux pump *qacEdelta1*, encoding a quaternary ammonium compound (QAC) efflux pump, was detected in eight Heidelberg chicken outbreak isolates (FML-M2-0001 through FML-M2-0004, FML-M2-0006, FML-M2-0007, FML-M2-0009, and FML-M2-0010). This gene has been linked to tolerance to QACs, and it may be selected for in environments where QAC-based sanitizers are used [51, 52]. The *asr* gene (anaerobic sulfite reduction), encoding an acid shock protein for survival in acidic conditions (such as acid-based antimicrobials [53]), was present in all but two NOA isolates (Enteritidis FML-M2-0028, NOA, raw chicken breast; Typhimurium var. 5-FML-M2-0029, NOA, chicken breast).

Higher resistance levels in some OA strains have been previously reported; Etter et al. found 1-10 AMR genes among seven Heidelberg strains from the 2013-2014 chicken outbreak [23], while phenotypic testing of 2015 roast pork outbreak monophasic Typhimurium strains showed most were multi-drug resistant [54]. Conversely, Procura et al. found few AMR genes in *S. enterica* from chicken livers, except erythromycin resistance in all isolates and streptomycin resistance in 22% [55]. In a study comparing prevalence of AMR in animal and nut products, the authors found most *S. enterica* from raw almonds (n =73/83) were resistant to two or fewer of 15 antimicrobials, with 52 being pan susceptible [56].

#### Heavy Metal Tolerance Genes

Heavy metal carriage has been previously linked with AMR gene carriage in livestock and human salmonellosis and may be carried on plasmids or other transferable elements [57–60]. Consequently, we assessed carriage of heavy metal resistance genes in our isolates. Carriage of genes conferring gold and arsenic tolerance was nearly ubiquitous among *S. enterica* isolates (**Figure 1**). All contained *golST* for gold tolerance, and all but two (Enteritidis FML-M2-0028, NOA, chicken breast; Typhimurium var. 5-FML-M2-0029, NOA, chicken breast) carried *arsR* or *arsABCDR* for arsenic tolerance. Copper (*pcoABCDERS* or *pcoACDRS*) and silver (*silABCEFPRS*) genes were found in all monophasic Typhimurium isolates (FML-M2-0011 through FML-M2-0025) and five 2013-2014 chicken outbreak Heidelberg isolates (FML-M2-0001, FML-M2-0003, FML-M2-0006, FML-M2-0009, and FML-M2-0010). Tellurium tolerance (*terDWZ*) was present in these five Heidelberg isolates and monophasic Typhimurium isolate FML-M2-0025 (NOA, pork feet). Mercury tolerance (*merAPR*, *merABDPRT*, or *merACDEPRT*) was observed in three chicken outbreak Heidelberg isolates (FML-M2-0002, FML-M2-0004, and FML-M2-0007), all roast pork monophasic Typhimurium isolates, and one kosher broiled chicken liver outbreak Heidelberg isolate (FML-M2-0030). Mercury tolerance did not co-occur with tellurium, copper, or silver tolerance among Heidelberg isolates, nor with tellurium tolerance among monophasic Typhimurium isolates.

Overall, monophasic Typhimurium isolates carried the greatest number of heavy metal tolerance genes, consistent with previous research [61]. Acquisition of these genes facilitates bacterial survival and has been noted to contribute to outbreak severity [61]. Furthermore, the highest frequency of both heavy metal tolerance and AMR genes was observed in Heidelberg isolates from the chicken outbreak and monophasic Typhimurium isolates from the roast pork outbreak, which was previously reported for *S. enterica* from livestock, animal foods and carcass samples in the EU, where heavy metals are often used as growth promoters [57].

### Sanitizer Tolerance

#### Sodium hypochlorite (NaOCl)

Acquired tolerance to sanitizers used in food processing environments can contribute to survival and persistence of *S. enterica* [62], increasing the risk of potential outbreaks. Exposure to sublethal doses can lead to bacterial repair mechanisms and allow for subsequent survival in extreme conditions [62]. Sanitizer tolerances are shown in **Figure 1** with detailed values in **Table S2**. NaOCl MIC among isolates FML-M2-0001 through FML-M2-0048 averaged >200 ppm in 1X TSB and 113 ppm in 1/20X TSB, indicating that nutrient concentration influences MIC values (*p* <0.05). MIC varied by serovar (*p* <0.05), with serovar Typhimurium having the highest MIC (>200 ppm), followed by Enteritidis (125 ppm), Heidelberg (109 ppm), Newport (111 ppm), and monophasic Typhimurium and Typhimurium var. 5-(100 ppm). Typhimurium var. 5-isolates had a higher MIC than monophasic Typhimurium averaged across both conditions (*p_adj_* <0.05). Isolates from the roast pork (FML-M2-0011 through FML-M2-0023), chicken (FML-M2-0001 through FML-M2-0010), kosher broiled chicken liver (FML-M2-0030 through FML-M2-0032), raw almond (FML-M2-0033), and peanut butter (FML-M2-0047 through FML-M2-0048) outbreaks had mean NaOCl MIC values at 1/20X TSB of 92 ppm, 95 ppm, 158 ppm, 150 ppm, and >200 ppm, respectively, though differences were not significant (*p_adj_* >0.05). NOA isolates averaged 122 ppm in 1/20X TSB.

Previous studies have also reported high NaOCl tolerance among *S. enterica* isolates [63–66]. Xiao et al. found >256 ppm MIC when using 11.92% NaOCl for most poultry supply chain-derived isolates (n =161/172) [63], while Obe et al. observed MIC values of 500-1,000 ppm using 12.5% NaOCl [64]. Humayoun et al. reported a mean MIC of 3,152 ppm [65] with a lower active chlorine concentration (5.25-6.15% NaOCl). Sublethal NaOCl concentrations (50 ppm) were marginally more effective against Heidelberg than Typhimurium and Enteritidis isolates [67], consistent with this study.

Higher NaOCl tolerance has been linked to AMR and the *qacEdelta1* efflux pump [63]; however, Heidelberg isolates in this study carrying both (n =8; **Figure 1**) had a lower mean MIC than other isolates. The high and moderate MIC values of nut-associated isolates (>200 ppm in 1X, Typhimurium FML-M2-0047 and FML-M2-0048, OA, peanut butter; 150 ppm in 1X, Enteritidis FML-M2-0033, OA, almond) may be due to their low-moisture food matrices, which have been associated with cross-tolerance to NaOCl [68]. The concerningly high NaOCl tolerance observed in this study and others may contribute to *S. enterica* persistence in the food processing environments where this sanitizer is used.

#### Peracetic Acid (PAA)

PAA MIC values for isolates FML-M2-0001 through FML-M2-0029 averaged 176 ppm in 1X TSB and 26 ppm in 1/20X TSB; isolates FML-M2-0030 through FML-M2-0048 were not tested. In 1/20X TSB, all isolates had MIC values of 25 ppm, except for monophasic Typhimurium FML-M2-0011 (50 ppm; OA, roast pork). Nutrient concentration influenced MIC values (*p* <0.05), with nearly all isolates exhibiting higher MIC values in 1X TSB than 1/20X (*p_adj_* <0.05). MICs varied little by serovar or outbreak, remaining between 25-27 ppm in 1/20X and 170-200 ppm in 1X TSB. Monophasic Typhimurium had the lowest MIC in 1X (173 ppm), while Enteritidis and Typhimurium var. 5-had the highest (>200 ppm). However, serovar was not a significant determinant of MIC (*p* >0.05). By outbreak status, the lowest MIC in 1X TSB was among Heidelberg chicken outbreak isolates (170 ppm), while NOA isolates had the highest (183 ppm), though differences were not significant (*p_adj_* >0.05). Overall, PAA tolerance was lower than NaOCl, though nutrient concentration appeared more influential in PAA tolerance.

PAA tolerance findings aligned with previous studies. Etter et al. reported an average MIC of 73.6 ppm among MDR *Salmonella* Heidelberg isolates from the 2013-2014 chicken outbreak testing concentrations of 25-250 ppm (0.0025-0.025%) [23]. Mourao et al. found an MIC of 60-70 ppm in *S. enterica* from chicken meat using 5-90 mg/L PAA (0.0005-0.009%) [69]. In contrast, Jolivet-Gougeon et al. reported an MIC of 7 ppm PAA (0.0007%) for *Salmonella* Typhimurium LT2 [70]. Humayoun et al. found a much higher average MIC of 880 ppm in 88 MDR *S. enterica* isolates using PAA concentrations of 80-15,104 μg ml^-1^ (0.01-1.5%) [65], but found no association between MDR and increased PAA tolerance. Micciche et al. reported MICs of 500 ppm for household PAA (0.00625-0.4%), and 1,000 ppm for industrial grade PAA (0.00625-0.4%) in *Salmonella* Typhimurium derived from animals [71].

*Salmonella* Typhimurium can withstand potentially lethal acid shock (pH < 4.0) following adaptation to milder acidic conditions [72], which may partially explain the high MIC values among Typhimurium isolates. Acid tolerance can also be partially enhanced by pre-exposure to other stressors [73], which may be more prevalent in food processing environments [74]. This study did not evaluate PAA tolerance among non-food processing derived isolates (i.e. soil, water, and farm swabs: FML-M2-0036 through FML-M2-0038 and FML-M2-0043, FML-M2-0044, and FML-M2-0047), limiting evaluation of food processing stress effects on PAA tolerance. It is important to note that MIC values observed in all previous studies were below the 2,000 ppm PAA limit set by the USDA-FSIS [75], though advisable concentrations may vary by product. These findings emphasize the importance of using sanitizers at full working concentrations to avoid tolerance and avoid inducing antimicrobial resistance [76].

### Heat Shock

Development of heat tolerance can allow for bacterial survival of *S. enterica* through food processing and potentially incomplete cooking, as well as provide cross-protection to other stresses [24]. We had previously identified unusual heat tolerance in an outbreak-associated *S. enterica* serovar Heidelberg, including a strain associated with illness from ready-to-eat rotisserie chicken [23, 40]. Consequently, we investigated whether this might be a common strategy in our outbreak-associated isolates in this study. We assessed the heat tolerance of 11 OA isolates (FML-M2-0009 through FML-M2 0013, FML-M2-0030 through FML-M2-0033, FML-M2-0047 and FML-M2-0048) from four serovars at 56°C (**Figure 2**) after conducting growth curves (data not shown) to confirm comparability of growth profiles. All isolates survived at least 15 minutes of heat shock, 10 survived past 30 minutes, nine past 45 minutes, and three past 60 minutes. The most heat tolerant isolates (Heidelberg FML-M2-0010, OA, chicken and monophasic Typhimurium FML-M2-0012, OA, roast pork) had 30.6 CFU/mL and 575.0 CFU/mL, respectively, after 60 minutes. Serovar (*p* <0.05) and time point (*p* <0.05) influenced heat tolerance, but outbreak status did not (*p* >0.05). Bacterial counts (CFU/mL) did not significantly decrease after 3 minutes scald (*p_adj_* >0.05) but decreased after 6 minutes (*p_adj_* <0.05). Enteritidis isolate (FML M2-0033, OA, raw almonds) was more heat tolerant than monophasic Typhimurium (FML-M2-0011 through FML-M2-0013, OA, roast pork; *p_adj_* <0.05) and Typhimurium isolates (FML M2-0047 and FML M2-0048, OA, peanut butter; *p_adj_* <0.05).

**Figure 2:**
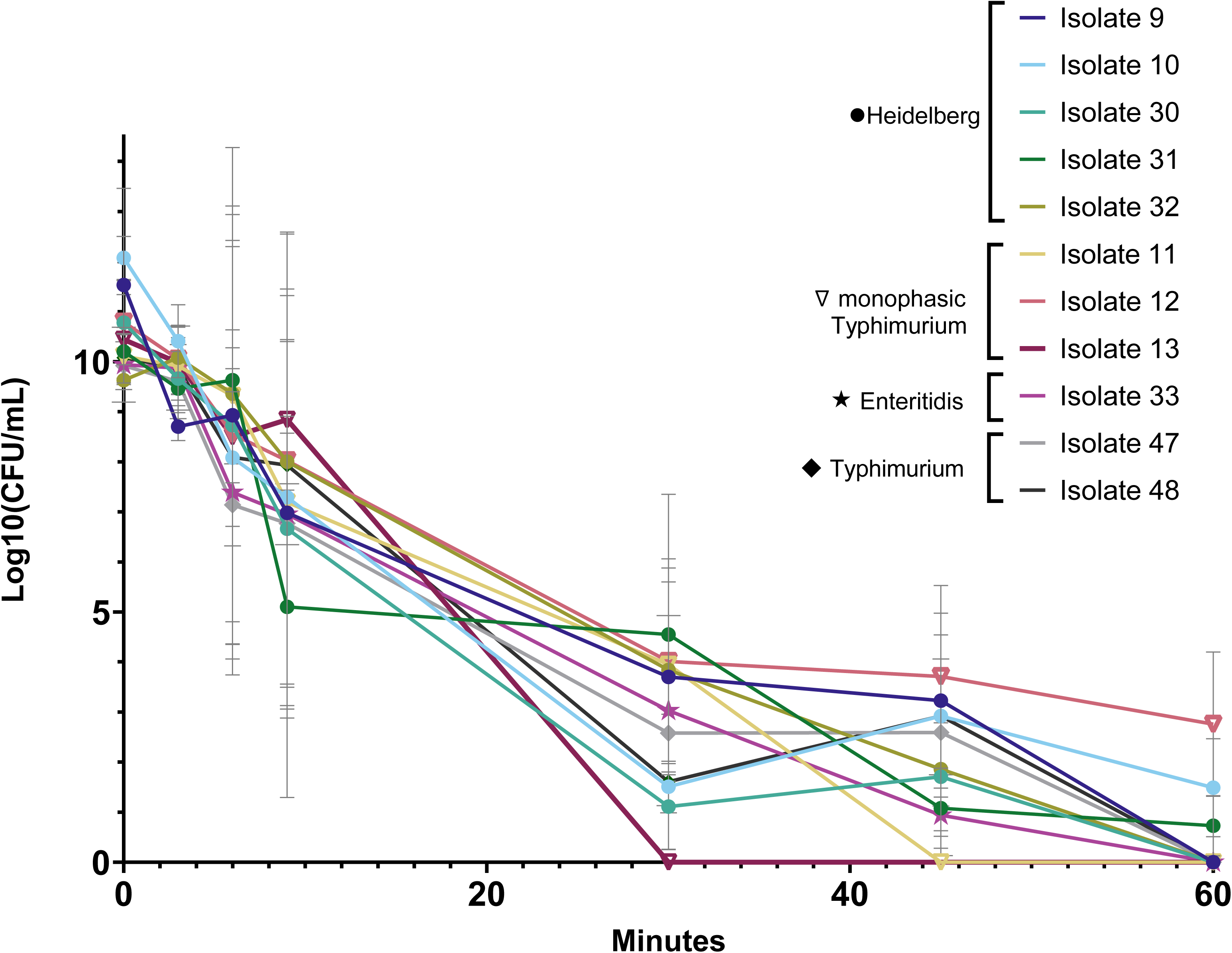
Heat tolerance of isolates (n=11) in stationary phase undergoing heat shock at 56°C. Time for isolates to reach 56°C from 37°C incubation (i.e. come-up time) was 2 minutes, 21 seconds.

Survival of poultry-associated strains at 56°C is concerning, where scalds in poultry processing typically last <2 minutes at 51-54°C for soft scalds and 59-64°C for hard scalds [14]. Three isolates from the 2013-2014 chicken outbreak (FML-M2-0002/R1-002, FML-M2-0005/R1-0004, FML-M2-0008/R1-0007) previously displayed enhanced heat tolerance [23], whereas only FML-M2-0010 did in the present study. However, all strains survived beyond typical scald durations. Inadequate cooking, as noted in the kosher broiled liver outbreak (FML-M2-0030 through FML-M2-0032 in **Figure 2**) [39], and possibly the roast pork (FML-M2-0011 through FML-M2-0013) and chicken outbreaks (FML-M2-0009 and FML-M2-0010), may have contributed to survival. Dawoud et al. reported that sublethal heat exposure can enhance bacterial thermal resistance, suggesting that prior heat exposure during food processing or cooking may have contributed to the survival observed in these outbreaks [22]. Our *in vitro* heat shock assay supports that inherent heat tolerance may have played a role.

Burns et al. found comparable heat tolerance in monophasic Typhimurium isolates from pig feed production, with one highly heat tolerant strain with the ASSuT AMR profile (resistances to ampicillin (A), streptomycin (S), sulfisoxazole (Su), and tetracycline (T)); isolates with the same AMR profile were implicated in the roast pork outbreak [54, 77]. However, only two of our three heat tolerant isolates were MDR, and prior research found no consistent AMR-heat tolerance association among poultry outbreak Heidelberg isolates [23].

While the heat tolerance of peanut butter and raw almond isolates was high, this temperature is substantially lower than that used in nut processing (>95°C) [16–18]. Other studies have demonstrated *S. enterica* survival in peanut butter at temperatures up to 120°C [78], and thermal resistance in peanut butter is well documented [79]. Ma et al. found that *Salmonella* Tennessee outbreak strains survived for 50 minutes at 90°C, with a 1-log CFU/g reduction, and were only undetectable after 120 minutes [80]. Similarly, Shachar and Yaron observed that peanut butter derived isolates of *Salmonella* Agona, Enteritidis and Typhimurium only had a 3.2-log reduction after 50 minutes at 90°C, with even lower efficacy at 80°C and 70°C [81]. Furthermore, *Salmonella* in dry products, such as nuts, are more heat resistant [82]. We found contradictory thermal resistance among our isolates from dry products; the raw almond isolate FML-M2-0033 was more tolerant compared to peanut butter isolates FML-M2-0047 and FML-M2-0048 (*p* <0.05). Regulations regarding almond processing changed following the almond outbreaks to require a minimum of 4-log CFU reduction of *Salmonella* from heat treatments [83], necessitated following large influxes of illness.

Previous studies have observed variation in heat tolerance by serovar and strain, similar to our findings. Two studies reported that *Salmonella* Enteritidis isolates were generally more resistant than Typhimurium [6, 82], while another study did not find differences by serovar substantial [6]. Hosts with higher internal body temperatures (e.g., chicken: 42°C) may activate thermal stress resistance mechanisms [22], which may explain enhanced heat tolerance among poultry-derived Heidelberg. Enhanced survival has also been observed in human illness-associated Enteritidis PT4 strains [84]. Lastly, we found potential cross-tolerances between heat stress and other environmental stresses. Induction of heat tolerance is known to provide subsequent cross-resistance to other stresses [24, 72], and we found that heat tolerant Heidelberg isolate FML-M2-0031 (OA, kosher broiled chicken liver) survived past 60 minutes of scald at 56°C, attached well in 1/20X TSB at 22°C, and also had an NaOCl MIC ≥ 200ppm. Heidelberg isolate FML-M2-0010 (OA, chicken) survived past 60 minutes of heat shock and attached strongly in 1X TSB at 22°C, but did not display additional phenotypic stress tolerances. Monophasic Typhimurium isolate FML-M2-0012 (OA, roast pork) had the highest CFU/mL after 60 minutes of heat shock and displayed enhanced attachment capacity but had relatively low NaOCl tolerance (50 ppm in 1/20X, 200 ppm in 1X).

Overall, isolates’ enhanced heat tolerance presents a concern for food processing facilities. According to the USDA-FSIS *Salmonella* Framework for Raw Poultry Products, a product is considered adulterated if it contains 10 CFU/mL or more of a serovar of public health concern [85]. After 60 minutes of a 56°C scald, significantly longer than used in processing for most products, we found two isolates were still above this level of contamination (Heidelberg FML-M2-0010, OA, chicken and monophasic Typhimurium FML-M2-0012, OA, roast pork), which is extremely concerning.

### Biofilm Attachment Assays

As the majority (80%) of bacterial infections in the U.S. are linked to foodborne pathogens residing in biofilms [31] and biofilms have been involved in several foodborne outbreaks [86–89], understanding attachment and biofilm formation capacity is crucial to identifying potential attributes to outbreaks. We evaluated all isolates at 24 h, 72 h, and 120 h for biomass (OD_600_) as an indication of attachment capacity and biofilm formation under four conditions: 1/20X 4°C (nutrient depletion, refrigeration); 1/20X RT (22°C) (nutrient depletion, room temperature); 1X 4°C (nutrient abundance, refrigeration); and 1X RT (22°C) (nutrient depletion, room temperature) (**Figure 3**).

**Figure 3:**
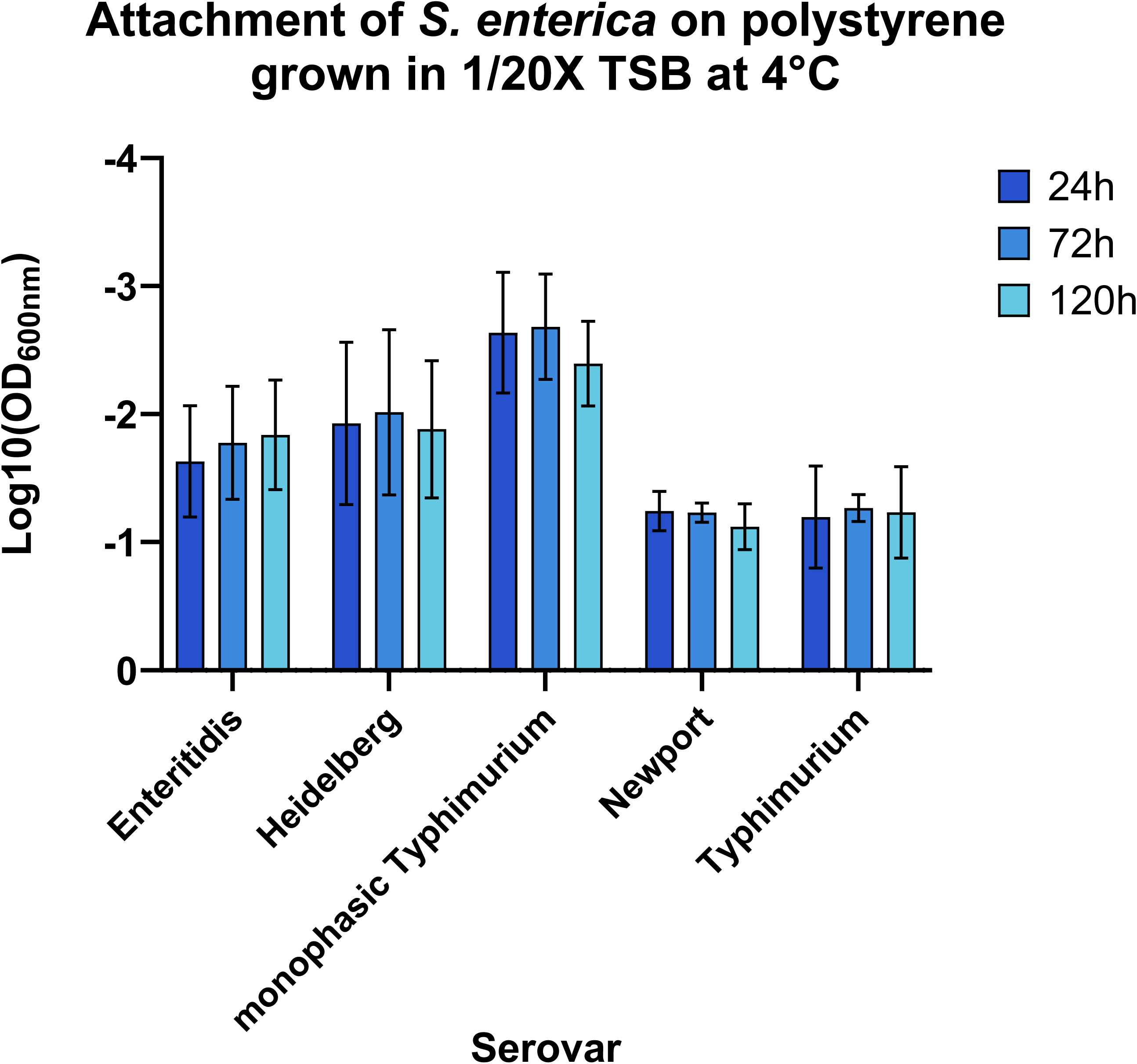
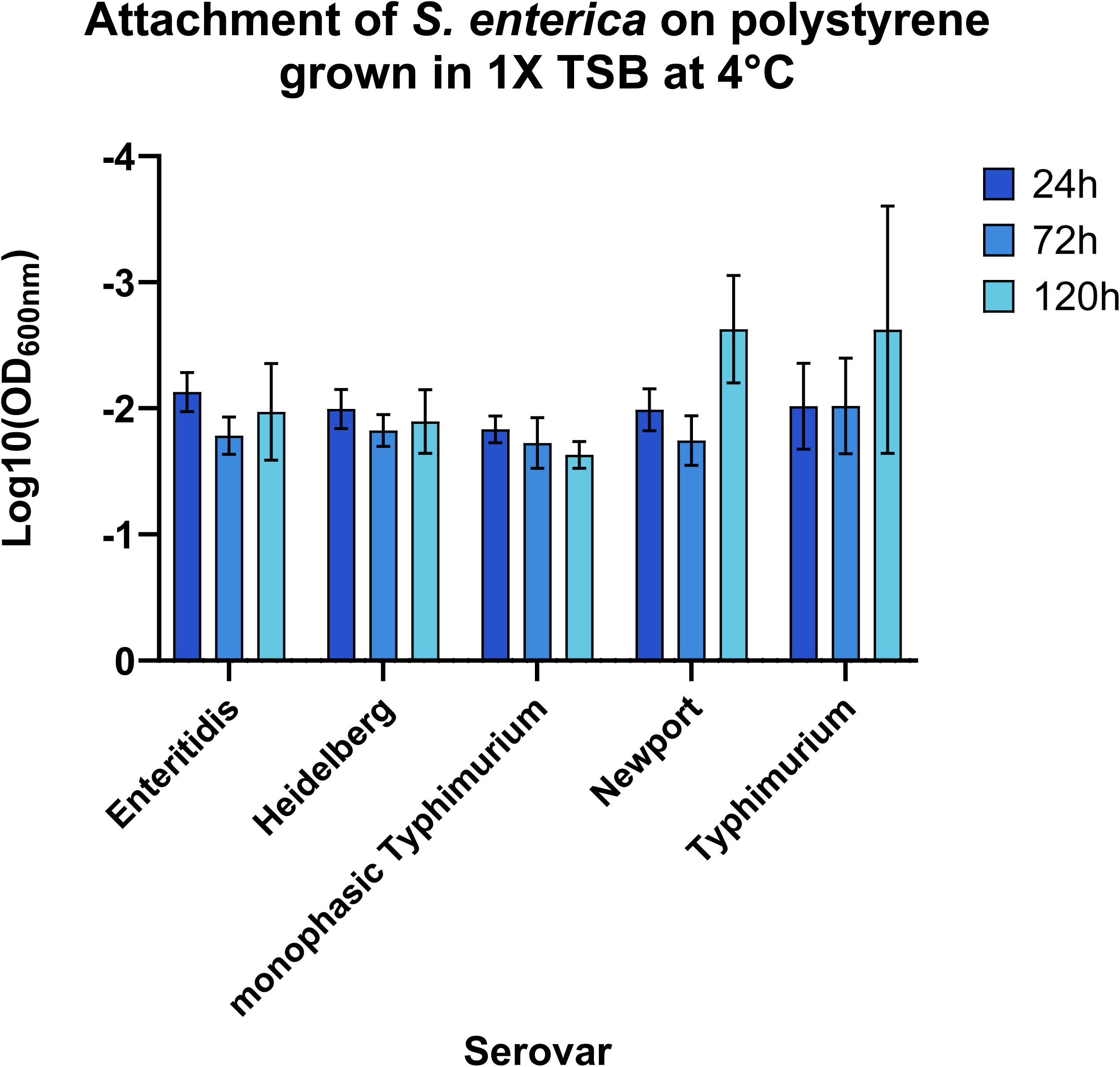
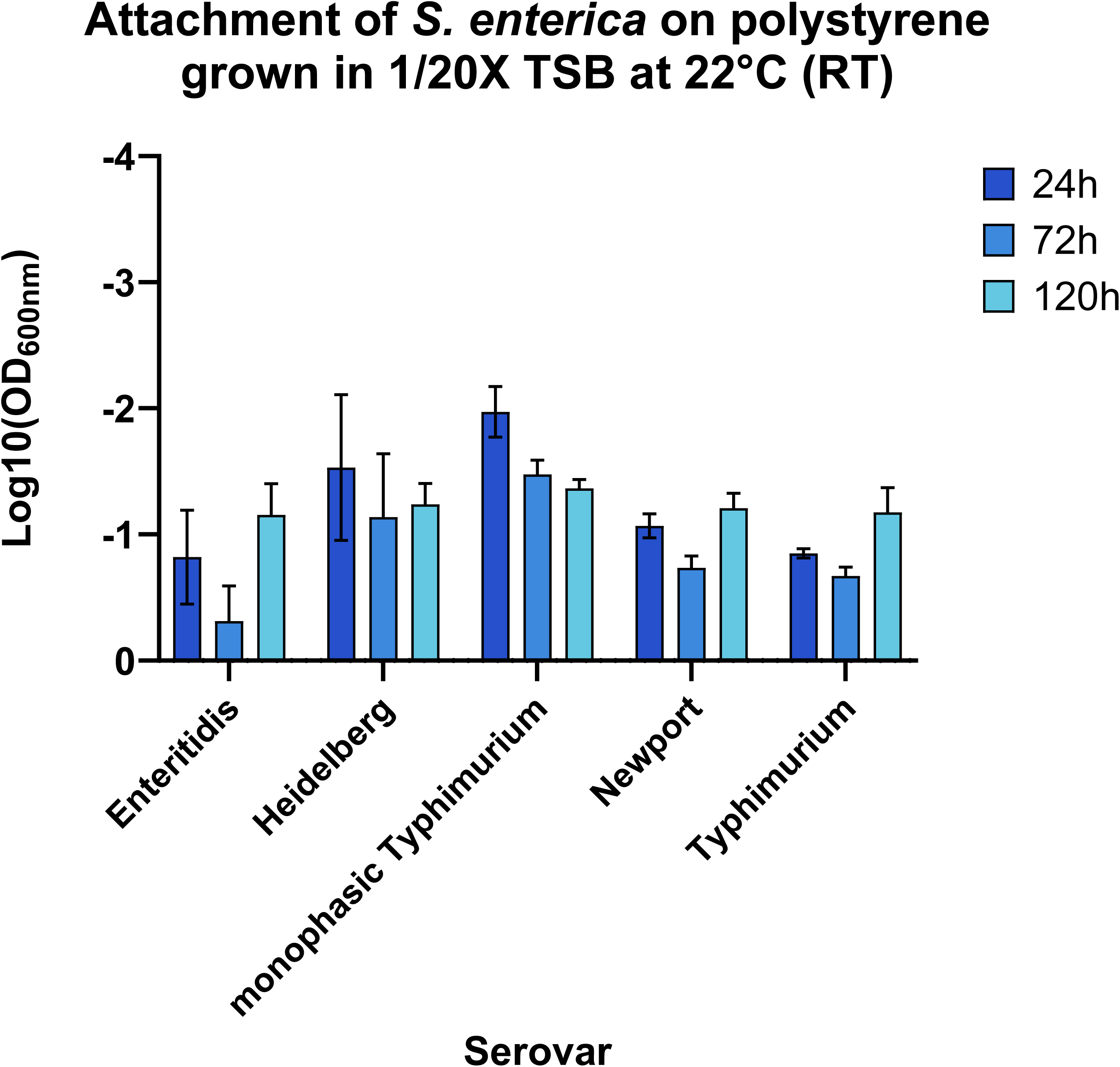
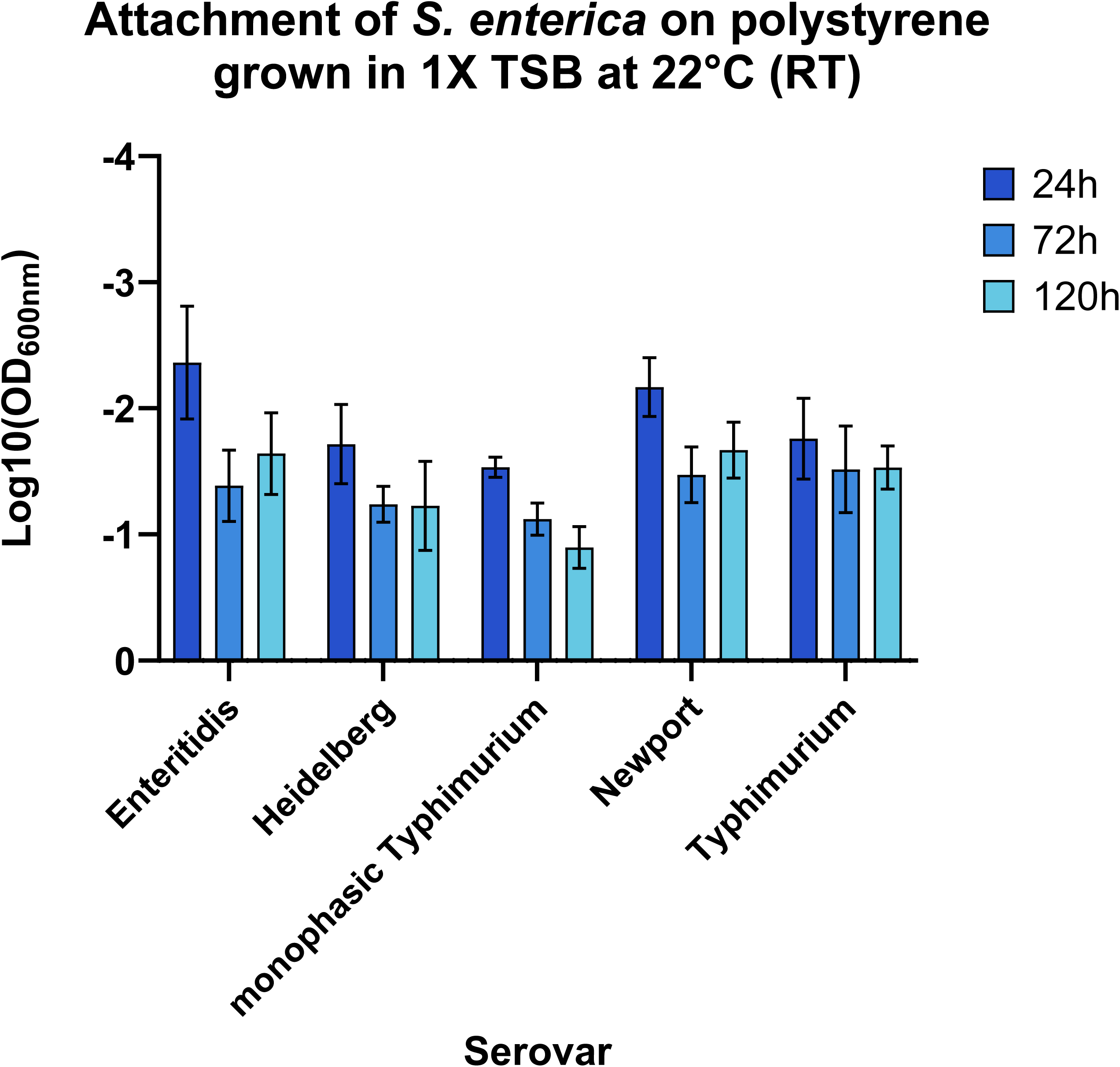
Attachment capacity of isolates, grouped by serovar, on polystyrene plates measured by crystal violet at 24, 72, and 120 hours in (A) 1/20X 4°C; (B) 1X 4°C; (C) 1/20X RT; and (D) 1X RT conditions, where a Log10(OD_600_) value closer to zero (shorter bar) indicates higher biomass and thus better attachment capacity.

In 1/20X 4*°*C, biomass varied by serovar (*p* <0.05), with Newport, Typhimurium, and Enteritidis having the greatest biomass, and monophasic Typhimurium with the least (*p_adj_* <0.05). Isolates Heidelberg FML-M2-0031 (OA, kosher broiled chicken liver) and Typhimurium FML-M2-0048 (OA, peanut butter) had the highest biomass (*p_adj_*<0.05). Biomass was unaffected by time point (24, 72, or 120 hours) (*p* <0.05) (**Figure 3A**). In 1X TSB, 4°C, biomass varied by serovar, time point, isolate, and serovar-time interactions (*p* <0.05). Biomass peaked at 72 h for all serovars except monophasic Typhimurium, which peaked at 120 h, outperforming all other serovars (*p_adj_* <0.05). Monophasic Typhimurium isolates had the highest biomass compared to all other serovars over all time points (*p_adj_* <0.05), and Typhimurium FML-M2-0012 (OA, roast pork) had a particularly high biomass (*p_adj_* <0.05). NOA isolates had greater biomass at 72 h than OA isolates, but OA isolates had greater biomass at 24 and 120 h, though not significant (**Figure 3B**).

At 1/20X RT, biomass varied by serovar, outbreak, time, isolate number, and serovar-time interactions (*p* <0.05). Enteritidis had the highest biomass (*p_adj_* < 0.05), while monophasic Typhimurium had the lowest (*p_adj_* <0.05). Biomass was highest at 72 h (*p_adj_* <0.05). At 72 h, Enteritidis had particularly high biomass (*p_adj_* <0.05), whereas monophasic Typhimurium had particularly low biomass, but exceeded all other serovars at 120 h. NOA isolates tended to form stronger biofilms compared to OA isolates, though not significant (**Figure 3C**). In 1X TSB RT, biomass varied by serovar, outbreak, time point, isolate number, and serovar-time interactions (*p* <0.05). Biomass was highest at 120 h compared to 72 h (*p_adj_* <0.05). Monophasic Typhimurium had the highest biomass across all serovars (*p_adj_* <0.05). OA isolates had higher biomass at all time points compared to NOA isolates (*p* <0.05). Heidelberg isolates had higher biomass than Enteritidis, Newport, and Typhimurium (*p_adj_* <0.05). The three kosher broiled chicken liver outbreak Heidelberg isolates (FML-M2-0030 through FML-M2-0032) had lower biomass compared to chicken outbreak Heidelberg isolates (FML-M2-0001 through FML-M2-0010). Specifically, FML-M2-0003 outperformed eight other isolates (*p_adj_* <0.05) (**Figure 3D**).

Overall, biomass was highest in 1/20X RT, followed by 1X RT. By serovar, Typhimurium isolates had the highest mean biomass in all conditions, followed by Newport, Enteritidis, Heidelberg, and monophasic Typhimurium. Monophasic Typhimurium isolates had the lowest biomass in 1/20X TSB, but the highest in 1X TSB. Enteritidis isolates had particularly high biomass values in 1X RT. Serovars Newport and Typhimurium also had particularly high biomass values in the high stress conditions of 1/20X 4°C. At room temperature, biomass at 72 and 120 h exceeded 24 h, suggesting that initial attachment is less temperature-dependent and maturation of biofilms is enhanced at RT compared to 4°C. OA isolates had greater biomass in 1X, RT at all time points (*p_adj_* <0.05) and marginally greater biomass in 1X, 4°C at 24 and 120 h (p >0.05). NOA isolates had higher biomass in 1/20X, 4°C at all time points (*p_adj_* <0.05), marginally higher biomass in 1/20X RT at all time points, and marginally higher biomass in 1X 4°C at 72 h.

Consistent with our findings, Stepanović et al. found that *Salmonella* biofilm formation in 1/20X TSB was more effective than 1X TSB across 122 strains [90]*. Salmonella*’s ability to form biofilms in response to starvation stress [91] is concerning, as 1/20X TSB conditions mimic the food processing industry [90]. Additionally, Stepanović et al. found that isolate source does not impact biofilm forming ability [90], which contradicts our findings where differences in biomass in 1X TSB were influenced by outbreak (*p_adj_* <0.05), though serovar distribution may be confounding. Agarwal et al. tested 151 strains across 69 serovars and found that over half formed moderate or strong biofilms [92]. Enteritidis outperformed Typhimurium in 1X 4°C, consistent with our results, and the highest biomass was observed at 48 h and 72 h, respectively [92]. Interestingly, Burns et al., noted biofilm formation as a potential future direction of research in their study evaluating heat tolerance among monophasic Typhimurium isolates from pig feed (Burns et al., 2016). Our findings suggest that monophasic Typhimurium may take longer to form mature biofilms, which could have contributed to persistence observed in the 2015 roast pork outbreak, as biofilm formation is crucial for environmental survival [93]. Further investigation into swine associated monophasic Typhimurium biofilm capacity is warranted. Lastly, Wang et al. linked sanitizer tolerance to biofilm forming ability [94]. In this study, Typhimurium – the strongest biofilm formers across all conditions— consistently exhibited the highest NaOCl tolerance, while the one Typhimurium isolate tested (FML-M2-0029) had the highest PAA tolerance, along with Enteritidis isolates.

## Conclusions

We assessed 43 OA and NOA *S. enterica* isolates from various serovars for phenotypic stress tolerance to heat and sanitizers, and for attachment capacity under processing-relevant conditions (nutrient abundance vs. depletion, and room temperature vs. refrigeration). Isolates displayed varying patterns of stress tolerance, with some showing potential cross-tolerance to multiple stresses. The three meat and poultry associated outbreaks included at least one isolate with survival following 60 minutes scald, suggesting that heat tolerance may have contributed to these outbreaks, particularly in cases of improper cooking. The short heat treatment used in the meat industry, aimed at preserving quality, emphasizes the concern regarding heat tolerance among meat-associated isolates. Nut-associated isolates also exhibited enhanced stress tolerance: Typhimurium isolates showed high biomass and NaOCl tolerance in 1/20X, and the Enteritidis isolate from almonds had high biomass and moderate NaOCl tolerance in 1/20X, as well as the highest survival across all time points during heat shock. Overall, the characteristics of these isolates likely contributed to the scope and severity of their respective outbreaks, and further research regarding the complex and nuanced relationship between specific *S. enterica* strains, stress tolerance, and their impacts to human health is needed to accurately model risks.

## Supplementary Material

**Figure S1:** Phylogenetic tree of isolates included in this study.

**Table S1:** Phenotypic AMR profiles for isolates in this study (from NCBI/isolate metadata).

**Table S2:** Tolerance to sodium hypochlorite (NaOCl) and peracetic acid (PAA) as minimum inhibitory concentrations (MIC; ppm).

## Acknowledgements

We thank and acknowledge researchers at the NY State Agency of Agriculture, the USDA-ARS, FDA CFSAN, and the Cornell Food Safety Lab, who provided isolates for this research. We also thank and acknowledge Tom Ford from Ecolab for providing a sample of PAA-based sanitizer to test, and John Johnston of USDA for facilitating transfer of isolates from USDA-ARS-NRRL in Illinois. This work was supported by the USDA National Institute of Food and Agriculture (NIFA) Agriculture and Food Research Initiative Strengthening and New Investigator Food Safety and Defense Program, project award no. 2019-06903. Additional salary support was received from the USDA Hatch funding mechanism via the Vermont Agricultural Experiment Station and gift funds provided to the Department of Animal Sciences at the University of Vermont for C. Patch’s contributions. Any opinions, findings, conclusions, or recommendations expressed in this publication are those of the author(s) and should not be construed to represent any official USDA or U.S. Government determination or policy.

