## Supplementary figures and images for "Stress Tolerance of Multiple *Salmonella enterica* Strains Associated with Foodborne Outbreaks"

### Supplemental Figure 1

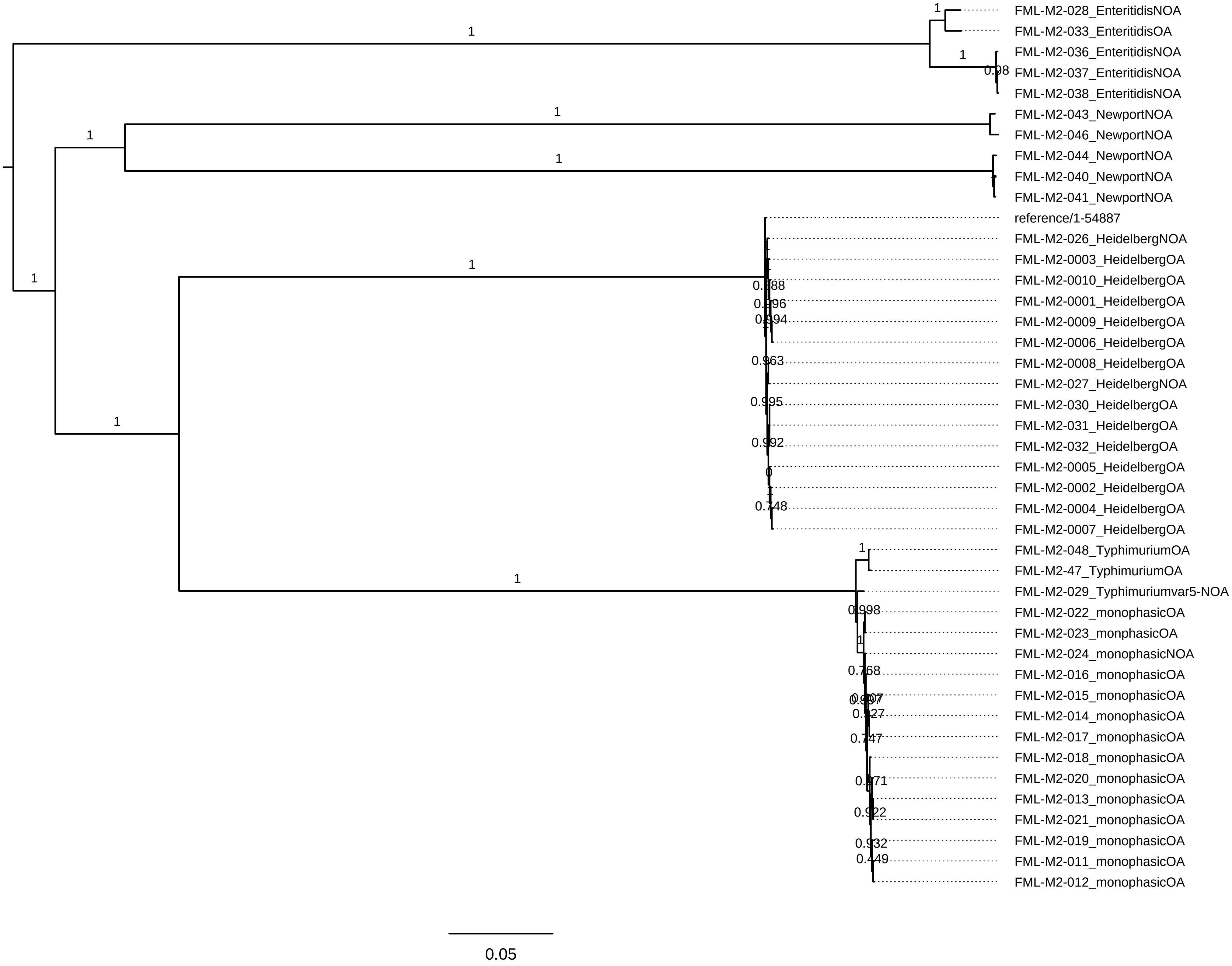
