## Supplemental Table 1 for "Stress Tolerance of Multiple *Salmonella enterica* Strains Associated with Foodborne Outbreaks"

**Table S1.** Phenotypic resistance profiles of isolates in this study.

| **Isolate Number** | **AMR phenotype †** |
| --- | --- |
| FML M2-0001 | GEN, KAN, STR, FIS, TET |
| FML M2-0002 | AMP, CHL, GEN, KAN, STR, FIS, TET |
| FML M2-0003 | GEN, KAN, STR, FIS, TET |
| FML M2-0004 | Unknown |
| FML M2-0005 | PAN-SUSC |
| FML M2-0006 | GEN, KAN, STR, FIS, TET |
| FML M2-0007 | Unknown |
| FML M2-0008 | PAN-SUSC |
| FML M2-0009 | Unknown |
| FML M2-0010 | Unknown |
| FML M2-0011 | Unknown |
| FML M2-0012 | Unknown |
| FML M2-0013 | Unknown |
| FML M2-0014 | Unknown |
| FML M2-0015 | Unknown |
| FML M2-0016 | Unknown |
| FML M2-0017 | Unknown |
| FML M2-0018 | Unknown |
| FML M2-0019 | Unknown |
| FML M2-0020 | Unknown |
| FML M2-0021 | Unknown |
| FML M2-0022 | [AMP, STR, FIS, TET ‡](https://www.ncbi.nlm.nih.gov/biosample/?term=SAMN04856237) |
| FML M2-0023 | [AMP, STR, FIS, TET ‡](https://www.ncbi.nlm.nih.gov/biosample/?term=SAMN04856237) |
| FML M2-0024 | [AMP, STR, FIS](https://www.ncbi.nlm.nih.gov/biosample/?term=SAMN04855045) |
| FML M2-0025 | [AMP, AXO, CIP (int)](https://www.ncbi.nlm.nih.gov/biosample/?term=SAMN05001758) |
| FML M2-0026 | [AMP, GEN, STR, TET](https://www.ncbi.nlm.nih.gov/biosample/?term=SAMN05201972) |
| FML M2-0027 | PAN-SUSC |
| FML M2-0028 | [AMC (int), AMP, STR, FIS, TET](https://www.ncbi.nlm.nih.gov/biosample/?term=SAMN05201730) |
| FML M2-0029 | [KAN, SUL, TET](https://www.ncbi.nlm.nih.gov/biosample/?term=SAMN05201583) |
| FML M2-0030 | Unknown |
| FML M2-0031 | Unknown |
| FML M2-0032 | Unknown |
| FML M2-0033 | Unknown |
| FML M2-0036 | Unknown |
| FML M2-0037 | Unknown |
| FML M2-0038 | Unknown |
| FML M2-0040 | Unknown |
| FML M2-0041 | Unknown |
| FML M2-0043 | Unknown |
| FML M2-0044 | Unknown |
| FML M2-0046 | Unknown |
| FML M2-0047 | Unknown |
| FML M2-0048 | Unknown |
| † AMC = amoxicillin-clavulanic acid, AMP = ampicillin, AXO = ceftriaxone, CHL = chloramphenicol, CIP = ciprofloxacin, FIS = sulfisoxazole, GEN = gentamicin, KAN = kanamycin, STR = streptomycin, SUL = sulfonamides, TET = tetracycline, PANSUSC = pansusceptible, Unknown = data not available  ‡ Indicates ASSuT resistance profile | |
