## Supplemental Table 2 for "Stress Tolerance of Multiple *Salmonella enterica* Strains Associated with Foodborne Outbreaks"

**Table S2.** Tolerance to sodium hypochlorite (NaOCl) and peracetic acid (PAA) sanitizers as minimum inhibitory concentrations (MIC; ppm).

|  |  |  | **Bleach MIC** | | **PAA MIC** | |
| --- | --- | --- | --- | --- | --- | --- |
| **Isolate Number** | **Serovar** | **Outbreak** | **1/20X** | **1X** | **1/20X** | **1X** |
| FSL M2-0001 | Heidelberg | chicken, 2013-2014 | 100 | >200 | 25 | >200 |
| FSL M2-0002 | Heidelberg | chicken, 2013-2015 | 50 | >200 | 25 | 100 |
| FSL M2-0003 | Heidelberg | chicken, 2013-2016 | 100 | >200 | 25 | >200 |
| FSL M2-0004 | Heidelberg | chicken, 2013-2017 | 100 | >200 | 25 | 100 |
| FSL M2-0005 | Heidelberg | chicken, 2013-2018 | 50 | >200 | 25 | >200 |
| FSL M2-0006 | Heidelberg | chicken, 2013-2019 | 100 | >200 | 25 | >200 |
| FSL M2-0007 | Heidelberg | chicken, 2013-2020 | 50 | >200 | 25 | >200 |
| FSL M2-0008 | Heidelberg | chicken, 2013-2021 | 100 | >200 | 25 | >200 |
| FSL M2-0009 | Heidelberg | chicken, 2013-2022 | 100 | >200 | 25 | >200 |
| FSL M2-0010 | Heidelberg | chicken, 2013-2023 | 100 | >200 | 25 | 100 |
| FSL M2-0011 | I 4,5,[12]:i:- | roast pork outbreak, 2015 | >200 | >200 | 50 | 100 |
| FSL M2-0012 | I 4,5,[12]:i:- | roast pork outbreak, 2016 | 50 | >200 | 25 | >200 |
| FSL M2-0013 | I 4,5,[12]:i:- | roast pork outbreak, 2017 | 50 | >200 | 25 | 100 |
| FSL M2-0014 | I 4,5,[12]:i:- | roast pork outbreak, 2018 | 100 | >200 | 25 | >200 |
| FSL M2-0015 | I 4,5,[12]:i:- | roast pork outbreak, 2019 | 100 | >200 | 25 | >200 |
| FSL M2-0016 | I 4,5,[12]:i:- | roast pork outbreak, 2020 | 50 | >200 | 25 | >200 |
| FSL M2-0017 | I 4,5,[12]:i:- | roast pork outbreak, 2021 | 50 | >200 | 25 | 100 |
| FSL M2-0018 | I 4,5,[12]:i:- | roast pork outbreak, 2022 | 100 | >200 | 25 | >200 |
| FSL M2-0019 | I 4,5,[12]:i:- | roast pork outbreak, 2023 | 50 | >200 | 25 | >200 |
| FSL M2-0020 | I 4,5,[12]:i:- | roast pork outbreak, 2024 | 50 | >200 | 25 | >200 |
| FSL M2-0021 | I 4,5,[12]:i:- | roast pork outbreak, 2025 | 100 | >200 | 25 | >200 |
| FSL M2-0022 | I 4,5,[12]:i:- | outbreak | 100 | >200 | 25 | >200 |
| FSL M2-0023 | I 4,5,[12]:i:- | outbreak | >200 | >200 | 25 | >200 |
| FSL M2-0024 | I 4,5,[12]:i:- | non-outbreak, 2016 | >200 | >200 | 25 | >200 |
| FSL M2-0025 | I 4,5,[12]:i:- | non-outbreak, 2016 | 100 | >200 | 25 | 100 |
| FSL M2-0026 | Heidelberg | non-outbreak, 2012 | 50 | >200 | 25 | >200 |
| FSL M2-0027 | Heidelberg | non-outbreak, 2007 | >200 | >200 | 25 | >200 |
| FSL M2-0028 | Enteritidis | non-outbreak, 2011 | >200 | >200 | 25 | >200 |
| FSL M2-0029 | Typhimurium var. 5- | non-outbreak, 2011 | 100 | >200 | 25 | >200 |
| FSL M2-0030 | Heidelberg | kosher broiled chicken liver, 2011 | 150 | >200 | n/a | n/a |
| FSL M2-0031 | Heidelberg | kosher broiled chicken liver, 2012 | >200 | >200 | n/a | n/a |
| FSL M2-0032 | Heidelberg | kosher broiled chicken liver, 2013 | 125 | >200 | n/a | n/a |
| FSL M2-0033 | Enteritidis | raw almonds, 2003-2004 | 150 | >200 | n/a | n/a |
| FSL M2-0036 | Enteritidis | non-outbreak, 2011 | 75 | >200 | n/a | n/a |
| FSL M2-0037 | Enteritidis | non-outbreak, 2011 | 125 | >200 | n/a | n/a |
| FSL M2-0038 | Enteritidis | non-outbreak, 2011 | 100 | >200 | n/a | n/a |
| FSL M2-0040 | Newport | non-outbreak | 150 | >200 | n/a | n/a |
| FSL M2-0041 | Newport | non-outbreak | 150 | >200 | n/a | n/a |
| FSL M2-0043 | Newport | non-outbreak | 100 | >200 | n/a | n/a |
| FSL M2-0044 | Newport | non-outbreak | 100 | >200 | n/a | n/a |
| FSL M2-0046 | Newport | non-outbreak | 100 | >200 | n/a | n/a |
| FSL M2-0047 | Typhimurium | peanut butter, 2009 | >200 | >200 | n/a | n/a |
| FSL M2-0048 | Typhimurium | peanut butter, 2010 | >200 | >200 | n/a | n/a |
